# MethCP: Differentially Methylated Region Detection with Change Point Models

**DOI:** 10.1101/265116

**Authors:** Boying Gong, Elizabeth Purdom

## Abstract

Whole-genome bisulfite sequencing (WGBS) provides a precise measure of methylation across the genome, yet presents a challenge in identifying regions that are differentially methylated (DMRs) between different conditions. A number of methods have been proposed which mainly focusing on the setting of two-group comparison. We develop a DMR detecting method `MethCP` for WGBS data, which is applicable for a wide range of experimental designs beyond the two-group comparisons, such as time-course data. `MethCP` identifies DMRs based on change point detection, which naturally segments the genome and provides region-level differential analysis. For simple two-group comparison, we show that our method outperforms developed methods in accurately detecting the complete DM region on a simulated dataset and an Arabidopsis dataset. Moreover, we show that `MethCP` is capable of detecting wide regions with small effect sizes, which can be common in some settings but existing techniques are poor in detecting such DMRs. We also demonstrate the use of `MethCP` for time-course data on another dataset following methylation throughout seed germination in Arabidopsis.

## 1 Introduction

DNA methylation is an important epigenetic mechanism for regulation of gene expression. Methylation is a process by which methyl groups are added to DNA cytosine (C) molecules. The methylation of promoter sites, in particular, is negatively correlated with gene expression while methylation in gene bodies is positively correlated with gene expression. Whole-genome bisulfite sequencing (WGBS) allows for precise measurement of DNA methylation across the genome. Briefly, when DNA is treated with bisulfite, the *unmethylated* cytosines are converted to uracil (U) leaving methylated cytosines unchanged. Sequencing of bisulfite-treated DNA and mapping of the sequenced reads to a reference genome then provides a quantification of the level of methylation at each cytosine. Methylation occurs in three different contexts: CG, CHG and CHH (where H corresponds to A, T or C). We will just refer to methylation of individual cytosine nucleotides.

In the analysis of BS-Seq data, a common interest is to identify regions of the genome where methylation patterns differ across populations of interest. Such regions are called differentially methylated regions (DMRs). Identifying DMRs are generally considered preferable than detection of individually differentially methylated cytosines (DMCs) from both statistical and biological perspective (Teschendorff and Relton, 2018). DNA methylation shows strong local patterns, and it is believed that region-level differences are more biologically important. Because of the low coverage and the fact that nearby cytosines usually have similar levels, combining them into regions substantially improves statistical power and lowers the false discovery. For the downstream analysis, reporting regions also reduces the redundancy.

A number of methods have been developed to identify regions from BS-Seq data that show differential methylation between two groups of samples (see Shafi *et al*. (2017) for a detailed review). One common strategy is to perform a test at every cytosine that appropriately accounts for the proportions and then use these significant results to determine the DMRs. For example, `methylKit` (Akalin *et al*., 2012) performs either a logistic regression test or Fisher’s exact test per cytosine; RADMeth (Dolzhenko and Smith, 2014) uses a beta-binomial regression, and a log-likelihood ratio test; `DSS` (Park and Wu, 2016; Feng *et al*., 2014; Wu *et al*., 2015) uses a Bayesian hierarchical model with beta-binomial distribution to model the proportions and tests for per-cytosine significance with a Wald test. Other methods use the local dependency between neighboring cytosines to improve their per-cytosine test. `BSmooth` (Hansen *et al*., 2012) and `BiSeq` (Hebestreit *et al*., 2013) both use local likelihood regression to estimate an underlying methylation curve, and then test for differences in the smoothed methylation ratios between populations. `HMM-DM` (Yu and Sun, 2016b,a) and `HMM-Fisher` (Sun and Yu, 2016; Yu and Sun, 2016a) both use Hidden Markov Models along the genome to account for the dependency between cytosines. For many of these methods, the region is often either predefined or determined by merging adjacent DMCs based on specific criteria such as distance.

Another approach is to directly segment the methylation levels to find DMRs. The method metilene (Jühling *et al*., 2016) uses a modified circular binary segmentation algorithm with statistics based on the mean differences in methylation ratios between two groups. The segments are tested for significance using Kolmogorov-Smirnov or Mann-Whitney U tests until the test results do not improve or the number of cytosines is too small.

We, too, propose a segmentation approach, `MethCP`, for finding DMRs from BS-Seq data. `MethCP` uses as input the results of a per-cytosine test statistic, like one of the methods described above, uses this input to segment the genome into regions, and then identifies which of those regions are DMRs. Our method, therefore, takes into account the coverages and biological variance between samples. Furthermore, all of the previously mentioned methods, including existing segmentation methods, are developed for simple two group comparisons, and are not straightforward to extend to more general experimental designs. `MethCP`, on the other hand, can be used in a wide variety of experimental designs.

We show via simulations that our method more accurately identifies regions differentially methylated between groups, as compared to competing methods. We illustrate the performance of `MethCP` on experimental data and show that its behavior on experimental data mirrors that of the simulations. We further demonstrate the flexibility of `MethCP` for use beyond the two-group setting by applying it to a time-course study.

## 2 Methods

`MethCP` assumes as input the results of a per-cytosine test of significance, such as those mentioned previously in the introduction. The main steps of `MethCP` are to 1) segment the test statistics into regions of similar values, and then 2) assign a p-value per region as to whether the region is a DMR. Let *T*_*k*_, *k* = 1, · · ·, *K*, be per-cytosine statistics for each of *K* cytosines, ordered by the location of the cytosines. We assume for now that the test statistics are independent (asymptotically) normally distributed, such as z-statistics or Wald statistics for testing equality of a proportion between two populations, and in 2.1 we extend this approach for other test statistics. We segment the *T*_*k*_ into regions of similar levels of significance based on the Circular Binary Segmentation (CBS) algorithm of Olshen *et al*. (2004), which was originally developed for segmentation of DNA copy number data. Note that `MethCP` applies the segmentation to test-statistics *T*_*k*_, which is a summary per cytosine as to the difference of interest across the samples, so that it finds regions of similar population significance.

Briefly, binary segmentation methods involve testing over all of the possible breakpoints (cytosines) for whether there is a change in the mean of *T* at location *i* ∈ [*K*]; in the case of genomic data, the segmentation is applied per chromosome. The CBS algorithm performs a binary segmentation and adapts the algorithm so as to view the data from a chromosome as if it lies on a circle, segmenting the circle into two arcs. The segmentation procedure of CBS is then as follows: for each possible arc defined by, 1 ≤ *i* < *j* ≤ *K*, the likelihood ratio test statistic *Z*_*ij*_ is calculated by comparing the mean value of *T*_*k*_ found in the arc from *i* + 1 to *j* with that found in the remaining circle. To find a significant breakpoint, CBS determines whether the statistic *Z =* max_1≤*i*<*j*≤*K*_ |*Z*_*ij*_| is significantly larger than 0. If so, this implies a detection that the arc (*i* + 1, *j*) has a significantly different mean than the remaining arc and the two arcs are declared to be separate segments. The procedure is then applied recursively on each resulting segment until no more significant segments are detected.

The number of computation required for the segmentation is *O*(*K*^2^). However, due to the uneven distribution of methylation cytosine across the genome, “gaps” where nearby methylation cytosines are far from each other and almost uncorrelated naturally presegment the genome. Like other methods (Jühling *et al*., 2016), `MethCP` can optionally presegment the genome and apply the algorithm separately to these highly separated segments. This reduces the computation to *O*(*KM*), where *M* ≪ *K* is the maximum number of cytosines in these segments.

The underlying purpose of the segmentation step is to segment the differential region from the undifferentiated regions. After the completion of the segmentation, it remains to determine which of these segmented regions correspond to significant DMRs, rather than their surrounding undifferentiated regions. To classify these regions, we use meta-analysis principles to aggregate the per-cytosine statistics and obtain one single statistic per region, to which we apply significance tests.

We assume that for each cytosine the calculated statistic *T*_*k*_ can be written as 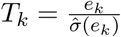, where *e*_*k*_ is the effect size that has approximate normal distribution with estimated variance 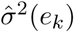. For a region *i* with a set of cytosines *S*_*i*_, the weighted effect size is given by

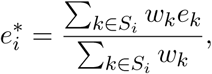

where the weights *w*_*k*_ signify the contribution of cytosine *k*. Typically in meta-analysis applications, *w*_*k*_ is set to be 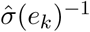(Borenstein *et al*., 2009). In the case of WGBS, assuming that appropriate methods which account for the variability in the counts are used to calculate *T*_*k*_, 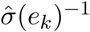 will be closely related to the coverage of the cytosine, which we designate as *C*_*k*_. Alternatively, for example, when 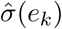 is not available, we can use *w*_*k*_ = *C*_*k*_, explicitly giving larger weights for high coverage cytosines.

A test statistic for a region found after segmentation is therefore calculated by 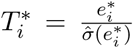, where 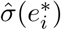 is the estimated variance of 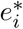. Based on our Gaussian distribution assumptions on the individual *e*_*k*_, we call the region significant if 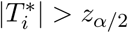, where *α* is the significance level.

In the standard meta-analysis, the individual *T*_*k*_ are often assumed to be independent so that the estimated variance of 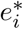 is given by

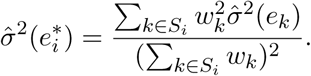

In the setting of methylation analysis, we have noted that the individual loci statistics are not independent. Even so, we show via simulation study (Section 3.1) that in using this estimate of 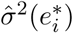, we still control the false discovery rate per region.

### 2.1 Generalizing beyond z-statistics

The above approach relies on input statistics that are Gaussian. This can be limiting, since methods often produce other types of statistics, such as Fisher’s exact test implemented by `methylKit` and log-likelihood ratio test from `RADMeth`. For this reason, we give a further adaptation in `MethCP` so as to be applicable for any cytosine-based parametric statistics that result in valid p-values. Let *p*_1_, *p*_2_, · · ·, *p*_*K*_ be the p-values indexed by the location of cytosines. For segmenting the genome into regions, we use the standard transform of the *p*-values to Z-scores,

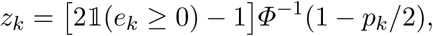

where *Φ* is the cumulative distribution function of standard Gaussian. `MethCP` then performs CBS on the *z*_*i*_’s to segment the genome.

Region-level statistics can be obtained by aggregating p-values using Fisher’s combined probability test (Fisher, 1934) or Stouffer’s weighted Z-method (Stouffer *et al*., 1949; Whitlock, 2005). Namely, for a region *i* with a set of cytosines *S*_*i*_, let

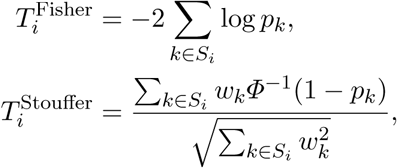

where *w*_*k*_ can be chosen to be constant or given by coverage *C*_*k*_. We test 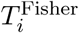 against 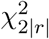, and 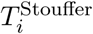 against standard Gaussian for significance.

### 2.2 Quantifying Region Alignment

To quantify the performance of the methods or similarity of DMR sets detected by different methods, we need to define some measures for whether a region was successfully detected. One simple solution is just to calculate measures of specificity and sensitivity based on the percentage of individual cytosines were correctly called to be in a DMR or not. However, since our goal is to detect *regions*, this is unsatisfactory. Thus, we extend the specificity and sensitivity to region detection problem. Our framework of evaluation is closely related to supervised measures such as directional Hamming distance and segmentation covering in the image segmentation literature (Huang and Dom, 1995; Pont-Tuset and Marques, 2016). Specifically, to determine whether a detected region is considered a true or false positive, we set a parameter *α* ∈ (0,1] that is the percentage of overlap required in order to be considered as having successfully detected a region. We then calculate true positive rates (TPR) and false positive rates (FPR) that vary with *α*.

Furthermore, we will see that some DMR methods are biased toward longer or shorter regions (Section 3), which can make comparing methods difficult. In order to account for different biases of regions found (in the following, we refer to number of cytosines in a region as the length of the region), we calculate the percent overlap between a detected region and a true region using three different denominators: that of the detected region, that of the true region and that of the union of the detected and true ones. The three measures can be interpreted as the local measures of precision, recall and Jaccard index. The first two allowed us to distinguish as to whether methods detected a high percentage of the true regions, versus if a high proportion of the detected regions were truly DMRs. The local Jaccard index allows us to measure the similarity between detected and true regions symmetrically.

We demonstrate our definitions using the local measure of precision – i.e., the overlap is determined by the proportion of the detected region that intersects the truth. Denote the detected and true region set as 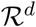 and 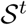, respectively. To determine whether a detected region 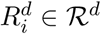 was a true positive (TP), we calculate the true positive indicator for 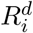:

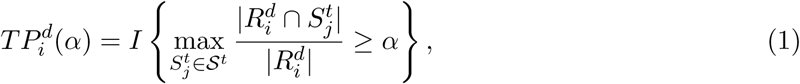

where 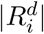 is the length of the detected region 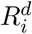, and 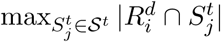 is the maximum overlapping cytosines of 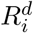 with a true region. Note that we take the maximum over all true regions to account for the fact that a detected region may overlap multiple true regions (and vice versa). From the 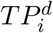 definitions, we calculate the total true positive (TP) as a function of 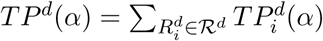.

The above formulas for *TP* can be extended to local measure of recall or Jaccard index by adjusting the denominator in Equation (1), from 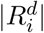 to 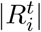 and 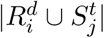, respectively, for calculating overlap. Similarly, we can calculate the false positive (FP), false negative (FN) and true negatives(TN). Please refer to Appendix C.1 for detailed formulations.

## 3 Results

### 3.1 Simulation Study

In this section, we applied our method as well as five representative methods (Shafi *et al*., 2017) `BSmooth`, `HMM-Fisher`, `DSS`, `methylKit` and `metilene` on simulated data with two population groups. Our method, `MethCP`, was run using the statistics of both `DSS` and `methylKit` as input (hereinafter referred to as `MethCP-DSS` and `MethCP-methylKit`). We evaluated their performances with the measures described in Section 2.2. The data simulation procedure and details of applying these methods can be found in Appendix B.

*A Simulation for Comparing Two Population Groups* Figure 1 shows the summary of the length of the DMRs detected by the six methods using the default significance level (or test statistic thresholds); we also show the distribution of the lengths of true regions. `MethCP` and `metilene` gives the closest length distribution to that of the true regions. Although we shortened the smoothing window compared to the default, `BSmooth` and `DSS` detect much larger regions. In contrast, `HMM-Fisher` and `methylKit` both detect small, fragmented regions.

**Fig. 1:**
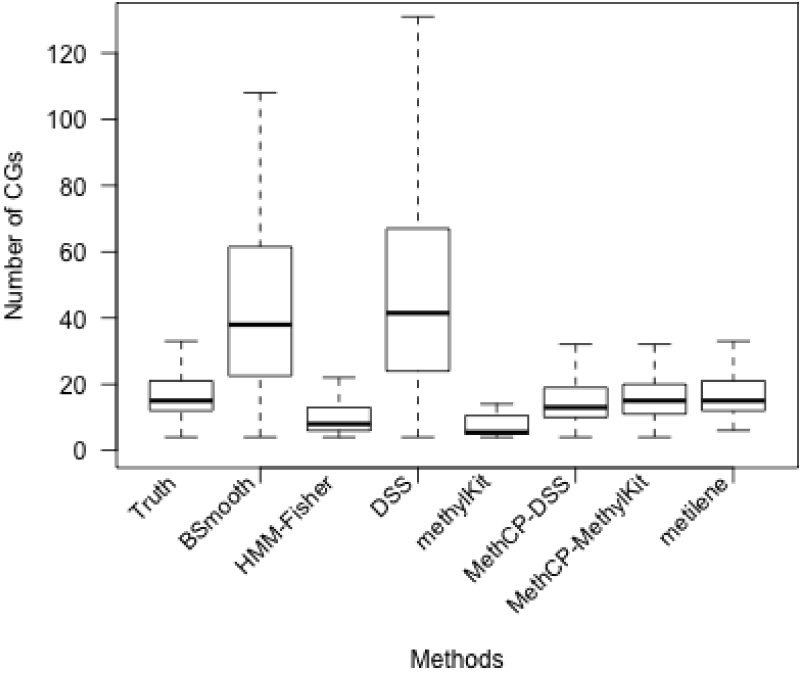
Boxplot of Number of CpGs in the DMRs. Number of CpGs in the true DMRs and in the DMRs detected by the seven methods compared for the simulated data.

To evaluate the accuracy of the methods on the simulated data, we plot the ROC curve for both the local precision (Figure 2a) and the local recall (Figure 2b). The local precision requires that a large percent of the detected region overlap a true DMR (easier for shorter detected regions and conservative methods), while the local recall requires that a large percent of the true region be overlapped by a detected region (easier for longer detected regions).

**Fig. 2:**
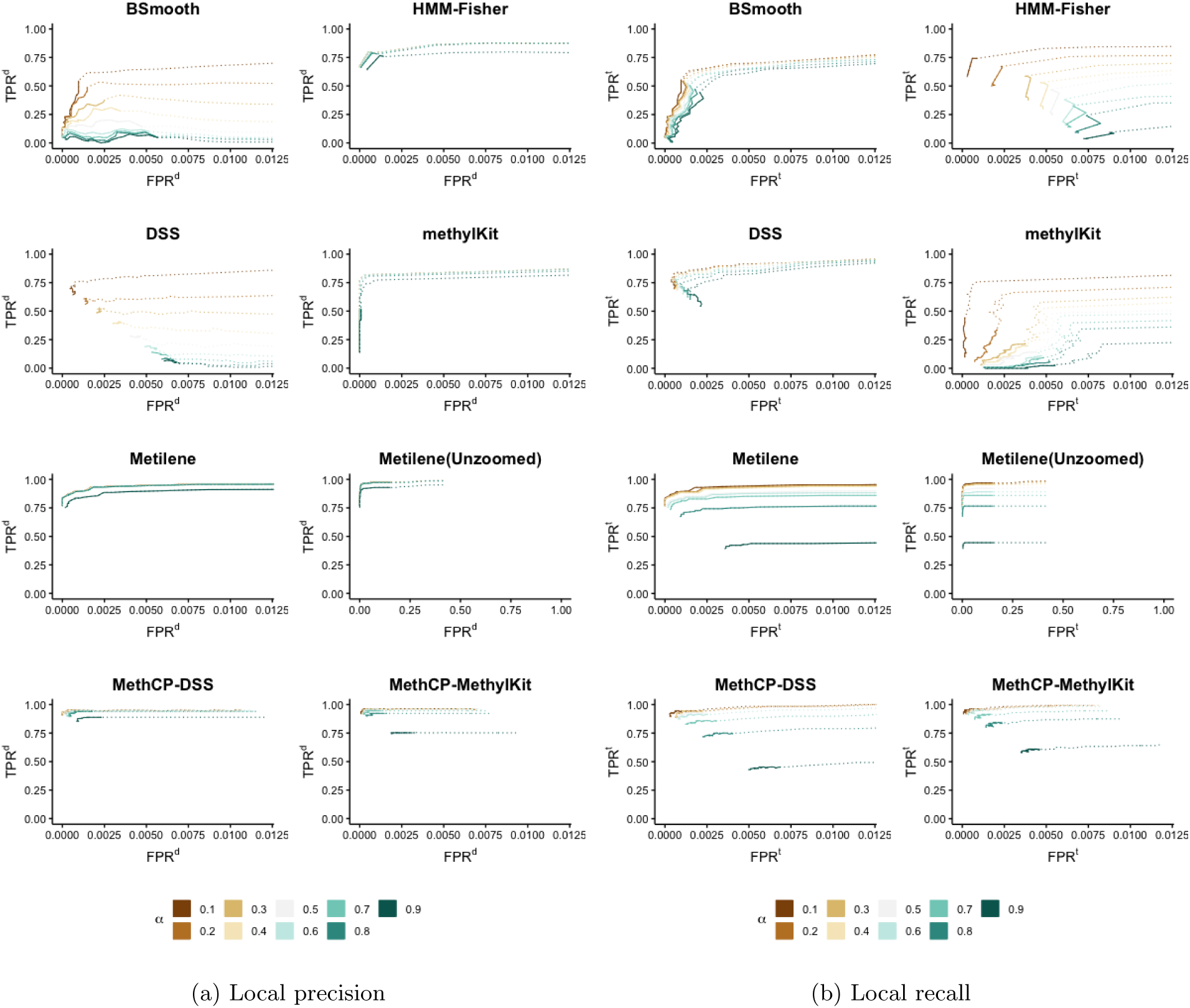
ROC curve: Method comparison on the simulated data. False positive rate (FPR) versus true positive rate (TPR) as we change the parameter *α* for (a) local precision measure and (b) local recall measure. The solid lines indicate significance levels smaller than 0.05 for `HMM-Fisher`, `DSS`, `metilene`, `MethCP-DSS` and `MethCP-methylKit`, statistics threshold larger than 4 for `BSmooth` (the author recommendation is 4.6), and *q*-values cutoffs less than 0.05 for `methylKit`. Thus, in real applications, we only focus on the regions of solid curves. For the completeness of the graph, we extend the curve to larger significance levels. `methylKit` uses an FDR correction procedure and only report *q*-values. Their false positive rate is reasonably controlled especially when we use the detected length as the denominator of our measure (FPR^*d*^).

`MethCP-DSS` and `MethCP-methylKit` detect highly similar regions despite using different test statistics as input. For small *α*, `MethCP` and `metilene` achieve the highest true positive rate across all of the methods. And both TPR^*d*^ and TPR^*t*^ are close to 1, which suggests that most true and detected regions match in pairs, with slight disagreement in the region border. This is not the case for other four methods, which for a given FPR are usually strong in either TPR^*d*^ or TPR^*t*^, but not both, which is evidence that either a proportion of the ground-truth regions are not detected (TPR^*d*^ high but TPR^*t*^ low), or a proportion of detected regions are not overlapping the ground-truth (TPR^*t*^ high but TPR^*d*^ low). `DSS` and `BSmooth` behave similarly in that TPR^*t*^ varies little with α while TPR^*d*^ decreases dramatically with the increase of *α*. This is an indication that both methods detect larger regions than the truth, which has been shown in Figure 1. Despite detecting regions wider than the true regions, `BSmooth` misses at least 20% of the true regions, as indicated by the values of TPR^*t*^, while `DSS` misses a smaller proportion.

`HMM-Fisher` and `methylKit` exhibit the opposite behavior, calling fewer and smaller regions significant, but as a result not obtaining good coverage of the true regions. The DMRs identified by these two methods are generally a subset of the true regions as indicated by the high values of TPR^*d*^ regardless of the significance level *α*. However, they also miss a good number of regions as shown in their lower TPR^*t*^ (Figure 2b). `metilene` and `MethCP` both rely on segmentation procedures, and `metilene` achieves better performance than the other competing methods, with high levels of both FPR^*d*^ and FPR^*t*^, though `metilene` still gives smaller TPR than `MethCP`.

However, in addition to assessing their overall sensitivity and specificity, we can consider whether the methods actually control the false positive rate at the desired level, and here we can see an even stronger difference between `metilene` and `MethCP`. In particular, for Figures 2a and 2b, the sold portions of the precision-recall curves indicate where the cutoff was due to the p-value cutoff being less than 0.05, meaning that control of the FPR at level 0.05 would be indicated by solid portion of the curve not extending beyond the true FPR of 0.05 given in the x-axis. For `metilene`, however, the solid curve continues beyond true FPR of 0.1 (see “Unzoomed” `metilene` plot), indicating that `metilene` is not accurately controlling the FPR. `MethCP`, on the other hand, does not have this property, and is quite conservative, despite having a higher TPR than the other methods.

*A Simulation for Small Effect Size Regions*. Recent studies (Breton *et al*., 2017; Leenen *et al*., 2016; Eichten and Springer, 2015) have shown that small-magnitude effect sizes are functionally important for DNA methylation and are associated with specific phenotypes. Furthermore, in plants, there exist other contexts of methylation other than CpG pairing (CHG and CHH methylation), where the baseline levels can be much lower, and hence their changes are also much smaller in scale, usually less than 10%. However, related discoveries have been hampered by a lack of DMR-calling tool addressing this issue (Breton *et al*., 2017; Eichten and Springer, 2015). We now show via a simulation study that `MethCP` is the only method of those we consider that is capable of accurately detecting regions with small (< 10%) changes in DNA methylation. In fact, none of the DMR detection methods we consider here, other than `metilene`, even identify *any* regions with methylation differences smaller than 10%, so we focus our comparison only on `metilene`. We simulated data sets with DMRs that have 2.5%, 5%, 10%, 20% changes in methylation, and their lengths vary from 25 methylation cytosines to 400 cytosines. As shown in Figure 3, `MethCP` achieves a high true positive rate while keeping the false positive rate low. `MethCP` consistently has higher TPR and lower FPR throughout the range of values.

**Fig. 3:**
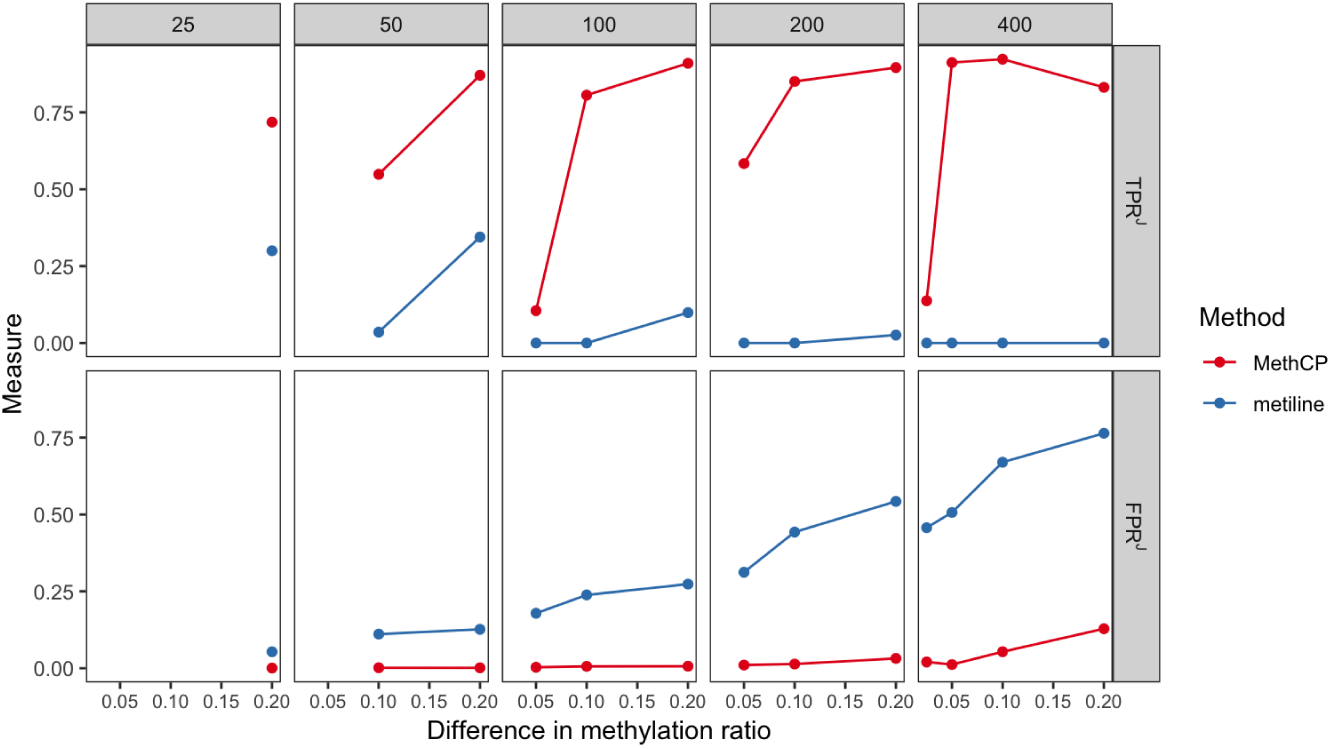
A comparison between metilene and MethCP for the small effect size simulation. Upper panel shows *TPR*^*J*^, and the lower panel shows *FPR*^*J*^ (*α* = 0.5). On the x-axis are the simulated methylation differences between control and treatment group. The five columns show simulated DMR length from short DMRs (25 cytosines) to large DMRs (400 cytosines).

### 3.2 Arabidopsis Dataset

To further illustrate the performance of `MethCP` on real datasets, we consider their performance on actual WGBS dataset. We do not have WGBS with actual ground truth (i.e., knowledge of where DMR regions are and are not located), so we use a dataset with a relatively large number of replicates that allow us to use permutation methods to give some idea of the performance of the tests; we combine this with a consideration of whether the important features seen in our simulation results are mimicked in real data. We use a WGBS on *Arabidopsis Thaliana* ((Coleman-Derr and Zilberman, 2012), GEO accession number GSE39045) which contained six biological replicates each of a wild-type control line and an H2A.Z mutant line.

Figure 4a shows the boxplot of the size of the DMRs detected (i.e., number of CpGs) under significance level 10^−2^ and FDR corrected level 10^−2^. We see that the sizes of the regions are mostly fairly consistent across the different significance levels, and that the relative sizes between methods resemble that of the simulated data. Namely, `DSS` and `BSmooth` detect large regions, `HMM-Fisher` and `methylKit` identified small ones, and `MethCP-DSS` and `MethCP-methylKit` identified highly similar regions despite different test statistics they use. `metilene`, stands out as sensitive to the significance level, with smaller required significance levels resulting in more small regions being reported (and `HMM-Fisher` finds no DMRs at more stringent significance levels).

**Fig. 4:**
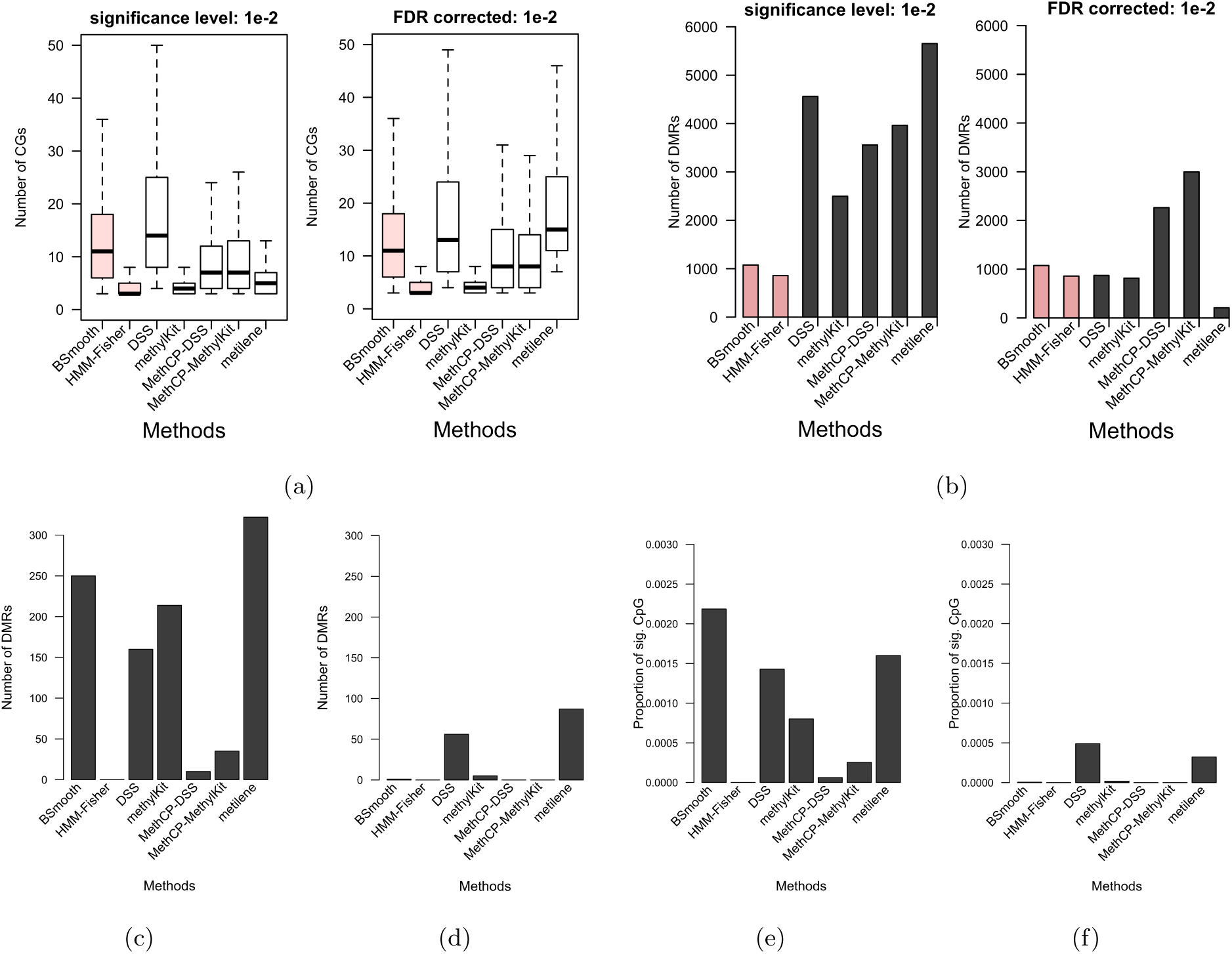
(a, b) Summary of DMRs detected for Arabidopsis dataset. (a) Boxplot of the number of CpGs in the DMRs. (b) Number of DMRs detected by different methods. We summarize the results under significance level 10^−2^ and FDR corrected level 10^−2^. `BSmooth` and `HMM-Fisher` are colored because we use the author recommended test statistic cutoff (4.6) for `BSmooth` and significance level 0.05 for `HMM-Fisher` on three plots (Small significance level for `HMM-Fisher` returns no DMR). **(c, d, e, f) Permutation results for Arabidopsis dataset**. (c) Permutation 1, number of DMRs detected. (d) Permutation 2, number of DMRs detected. (e) Permutation 1, the proportion of significant CpGs. (f) Permutation 2, the proportion of significant CpGs. Methods other than `BSmooth` uses significance level 0.01, and `BSmooth` uses the author recommended test statistic cutoff (4.6).

To compare the false positives produced by six methods, we create a null dataset, i.e., one with no region of cytosines different between the two groups, by randomly assigning the six controls into two groups of three. Individual differences between the mean methylation levels, combined with the fact that nearby sytosines have similar methylation levels in a sample, meaning that by chance there are regions of cytosines that are different between the two random groups. Therefore, we further permute the data in two different ways to create null data sets that have no DMRs between the two groups. The first method permutes the sequencing counts (methylated and unmethylated count pairs) across the samples for *each* CpG position (Permutation 1). The second method permutes the cytosine positions but keeps the sample labels, thus breaking any residual spatial signal between neighboring CpGs (Permutation 2). Figure 4 shows the number of DMRs and proportion of CpGs detected by each method. In both permutations, `MethCP` has a small number of DMRs relative to the other methods. Indeed for the second permutation, `MethCP` detects 0 DMRs. For the first permutation, only `HMM-Fisher` finds fewer, but `HMM-Fisher` also appears to have less power in detecting the real differences, Figure 4b. If we further consider how well `MethCP` assigns individual cytosines correctly to DMRs (i.e. its ability to detect of DMCs), then `MethCP` also results in fewer false assignments of individual cytosines than the other methods, which is surprising since it is not a DMC method.

We would note that despite detecting fewer false positive DMRs in our permutation analysis, `MethCP` still remains competitive, as compared to the other methods, in terms of the number of DMRs it finds on the real data (Figure 4b), indicating that it is not suffering from a lack of power. `DSS` and `methylKit` both find more regions, but we see from our permutation analysis that `DSS` and `methylKit` tend to find many more false positives than `MethCP`. Furthermore, the regions found by `methylKit` are small (Figure 4a), suggesting that like in the simulations `methylKit` may be missing or fragmenting large parts of the true DMRs.

### 3.3 Time-course dataset

A strength of `MethCP` is that it is flexible for a wide variety of experimental designs, because it only requires an appropriate per-cytosine test-statistic, which can be calculated by standard tests on a per-cytosine basis. We demonstrate this utility on a seed germination dataset from *Arabidopsis thaliana* (Kawakatsu *et al*. (2017), GEO accession number GSE94712). The data is generated from tracking over time germinating seeds after the dry seeds are given water. Two experimental conditions are considered: wild-type plants (*Col-0*) and *ros1*, *dml2*, and *dml3* triple demethylase mutants plants (*rdd*). ROS1, DML2 and DML3 are closely related DNA demethylation enzymes that mainly act in vegetative tissues. Two replicates were collected at each of 0-4 days after introduction of water (DAI), resulting in six time points, including the dry seed.

The original authors did not conduct a DMR analysis between these groups. Rather they did a two-group test of the overall methylation levels by grouping together the summarized methylation ratios and testing for the difference in distributions at each time point. They saw a modest overall increase between the two groups at each time point. We perform a DMR analysis with `MethCP` by fitting per CpG a linear model on the arcsine-transformed methylation ratios and choosing the statistic *T*_*k*_ to be the difference between the time coefficient of *Col-0* and *rdd* groups. Figure 6 shows the histogram of DMR length. Unlike the length distribution in Section 3.2, we detect some very large DMRs with small changes of methylation over time. Our ability to pick up both small and long regions gives us the ability to see multiple effects. We see the overall general increase, reported by the authors, represented by the long regions of small increase (Figure 5, regions colored black). But we also detect some smaller regions with an opposite pattern of decrease in methylation in the mutant samples (Figure 5, regions colored grey).

**Fig. 5:**
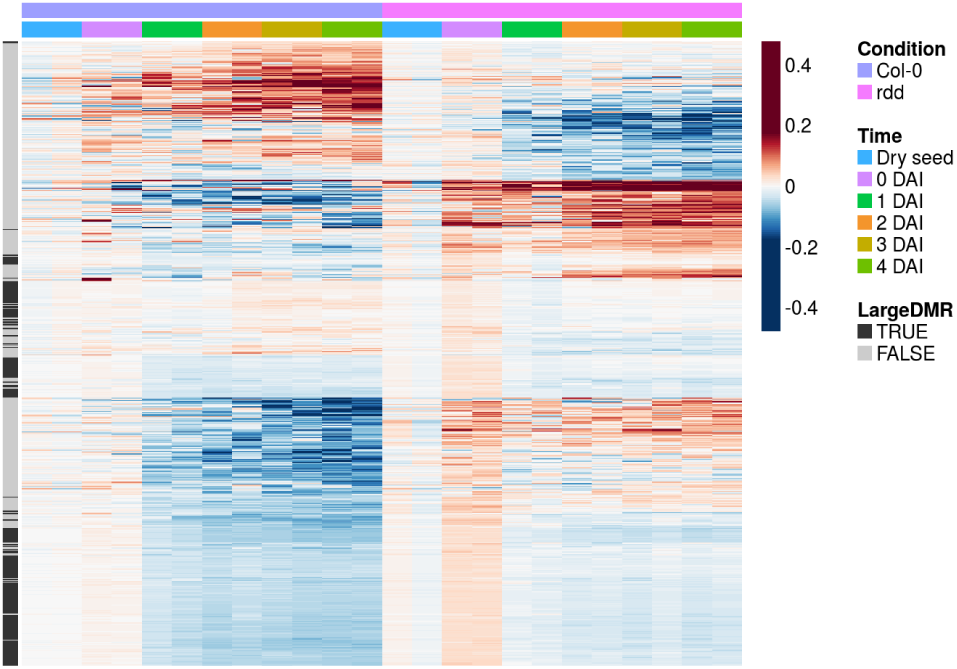
Heatmap of DMRs for the seed germination dataset. Heatmap of average methylation levels in the DMRs detected. Here we show the results for CpG context. For each condition and each DMR, we subtract the ratios by the average dry seed methylation ratios, so that the heatmap better shows the changes with time. We annotate the DMRs with greater than 200 CpGs as large regions.

**Fig. 6:**
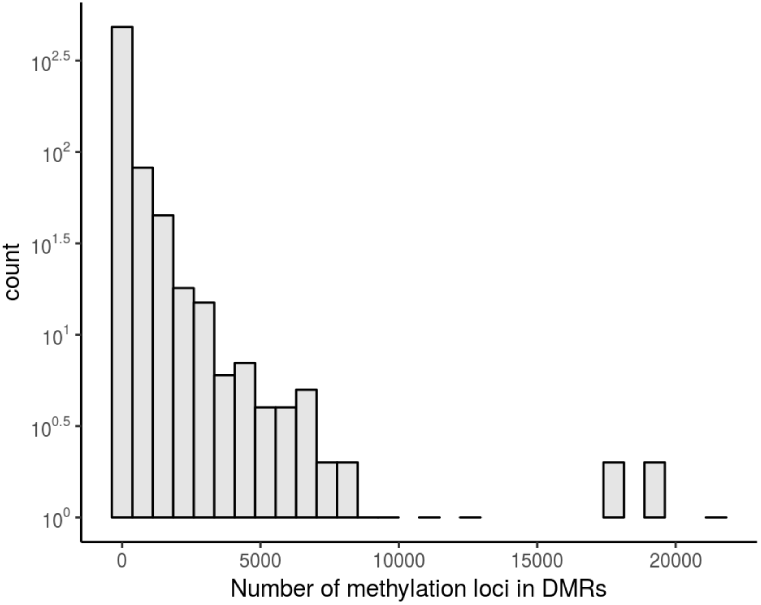
Lengths (number of methylation cytosine) of DMRs detected in the seed germination dataset (CpG context).

## 4 Conclusion

We proposed a method `MethCP` for identifying differentially methylated regions. We presented the results of `MethCP-methylKit` and `MethCP-DSS` on simulated and real datasets. And we showed that `MethCP` gives better accuracy and a lower number of false positives, as compared to existing methods. We also show that `MethCP` is the only method that detects large DMRs with small effect sizes, which is prevalent in DMR methylation data. Other than comparing two groups, we presented an example of a time-course study with `MethCP`. Our framework is in principle flexible for general experimental design assuming an appropriate single-cytosine test statistic can be calculated. Thus the method can be expanded immediately to more complicated situations, such as comparing multiple groups or measurements of methylation status over time or developmental progression.

## Funding

This work has been supported in part by a DOE BER grant, DE-SC0014081.

## A Quantifying Region Alignment: Measurement Formulations

Similar to 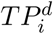 definitions, we calculate the total false positive (FP), and false negative (FN) as a function of *α*:

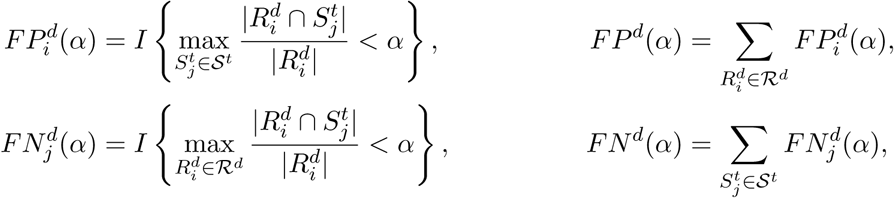

where the false negative is interpreted as the number of true regions that do have overlap greater than *α* with any detected positives.

The above formula can be extended to use local measure of recall or Jaccard index by adjusting the denominator in Equation 1, from 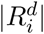 to 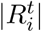 and 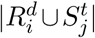, respectively, for calculating overlap.

For example, measuring the number of true positives, we have:

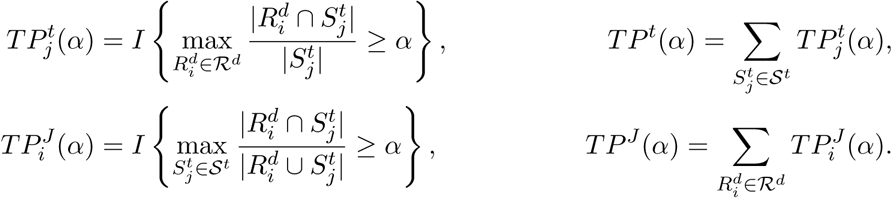

### Calculating True Negatives

Calculating the total number of true negatives would require a calculation of the number of detected regions that were truly not significant (i.e. equally methylated). However, an equally methylated region is a more nebulous quantity for a region (unlike for cy-tosines). Unlike the different DMRs, all equally methylated regions are equivalent from the point of view of all of these methods: arbitrarily defining separate regions within a large block of equally methylated regions could not be detected by any method. Instead, we use the following formula to get an estimation of the total number of true negatives:

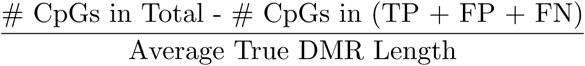

## B Simulation Study

### How we applied other methods

We applied our method as well as five representative methods (Shafi *et al*., 2017) `BSmooth`, `HMM-Fisher`, `DSS`, `methylKit` and `metilene` on both real and simulated data. Our method, `MethCP`, was run using the statistics of both `DSS` and `methylKit` as input. To be fair between methods, we remove the coverage filter for individual cytosines, as it varies by methods. Furthermore, the reads in symmetric CpG sites are collapsed. We set the same length filter (3 cytosines) and absolute mean methylation level difference filter (0.1) for DMRs, where the numbers are the default of the majority of methods. We shorten the smoothing window of `BSmooth` from default 1000 bps to 500 bps, which gives better results for our simulated dataset. For `DSS`, we use moving average smoothing, which is recommended in the documentation. For `methylKit`, the output is DMCs rather than DMRs. We combine adjacent DMCs to DMRs. Resulting DMRs that are smaller than 3 cytosines are discarded. All other parameters other than the significance level (test statistics cutoffs) were left at the default values.

### Generation of Simulated Data

We generate simulated BS-Seq data by the following procedure adapted from Yu and Sun (2016a). We assume there are *K* cytosines in the simulated genome and two groups of samples (“treatment” and “control”), each of size *n* = 3 to compare; this is similar to the level of replication that is often seen in WGBS. We designate regions within this genome to be classified as DMR by generating region size (number of CpGs) from a negative binomial distribution *NB*(*r* = 6, *p* = 0.25). We further require that the number of CpGs to be greater than 3 in each region. The starting positions of the DMR were chosen by random sampling. This divided the genome into differentially methylated and equally methylated regions.

To mimic the read coverage and the methylation ratio in real datasets, the actual sequencing counts were generated based on a human senescent cells dataset (Cruickshanks *et al*., 2013) available from the Gene Expression Omnibus (GEO) with accession number GSE48580. To determine the total read coverage, we randomly sampled from the observed coverage distribution of each cytosine in the human dataset. The number of reads determined to be methylated per cytosine was based on a binomial distribution, with the probability of proportion depending on what treatment group the sample was in and if the cytosine was in a DMR or not. For samples in the control group or for cytosines in the equally methylated regions, the binomial probability parameter was chosen from the observed distribution of the per-cytosine average methylation ratio in the senescent cells dataset. For DM regions, each DM region was randomly assigned one of five beta distributions from which the methylation probability of the treatment samples would follow; in addition, we require that the absolute difference between the mean of binomial distribution in treatment and in the corresponding control group is at least 0.2, which eliminated some of the five beta distributions from consideration. These beta distributions represent five different methylation levels, from poorly methylated to highly methylated (specific parameters of the beta distribution are given in Table 1). Then the cytosine methylation probability for samples in the treatment group was generated according to the beta distribution chosen for that DMR region. To take into account the high correlation of methylation levels between neighboring CpGs, we simulate smoothing DMR boundaries. For a DMR of length *l*, a region of length *w* ~ Unif(0.1*l*, 0.3*l*) is added to each side of the DMR where the methylation probability is given by a mixture of treatment and control. The weights of the treatment group decrease as we move to the edge of the DMR. In this paper, we show only results with the smoothing boundary, but simulation without smoothing boundaries give similar results.

**Table 1:** Parameters of beta distributions for simulating the methylated counts in the treatment group.

| Distribution | (a) | (b) | (c) | (d) | (e) |
| --- | --- | --- | --- | --- | --- |
| $\alpha$ | 2 | 6 | 10 | 14 | 18 |
| $\beta$ | 18 | 14 | 10 | 6 | 2 |
| Mean Probability | 0.1 | 0.3 | 0.5 | 0.7 | 0.9 |

### C Figures

**C.1 DMR lengths in the Time-course Dataset**

## References

Akalin, A., Kormaksson, M., Li, S., Garrett-Bakelman, F. E., Figueroa, M. E., Melnick, A., and Mason, C. E. 2012. methylkit: a comprehensive r package for the analysis of genome-wide dna methylation profiles. Genome biology 13, R87.

Borenstein, M., Hedges, L. V., Higgins, J., and Rothstein, H. R. 2009. Introduction to meta-analysis. Wiley Online Library.

Breton, C. V., Marsit, C. J., Faustman, E., Nadeau, K., Goodrich, J. M., Dolinoy, D. C., Herbstman, J., Holland, N., LaSalle, J. M., Schmidt, R., et al. 2017. Small-magnitude effect sizes in epigenetic end points are important in childrens environmental health studies: the childrens environmental health and disease prevention research centers epigenetics working group. Environmental health perspectives 125, 511.

Coleman-Derr, D. and Zilberman, D. 2012. Deposition of histone variant h2a. z within gene bodies regulates responsive genes. PLoS genetics 8, e1002988.

Cruickshanks, H. A., McBryan, T., Nelson, D. M., VanderKraats, N. D., Shah, P. P., Van Tuyn, J., Rai, T. S., Brock, C., Donahue, G., Dunican, D. S., et al. 2013. Senescent cells harbour features of the cancer epigenome. Nature cell biology 15, 1495.

Dolzhenko, E. and Smith, A. D. 2014. Using beta-binomial regression for high-precision differential methylation analysis in multifactor whole-genome bisulfite sequencing experiments. BMC bioinformatics 15, 215.

Eichten, S. R. and Springer, N. M. 2015. Minimal evidence for consistent changes in maize dna methylation patterns following environmental stress. Frontiers in plant science 6, 308.

Feng, H., Conneely, K. N., and Wu, H. 2014. A bayesian hierarchical model to detect differentially methylated loci from single nucleotide resolution sequencing data. Nucleic acids research 42, e69–e69.

Fisher, R. A. 1934. Statistical methods for research workers.

Hansen, K. D., Langmead, B., and Irizarry, R. A. 2012. Bsmooth: from whole genome bisulfite sequencing reads to differentially methylated regions. Genome biology 13, R83.

Hebestreit, K., Dugas, M., and Klein, H.-U. 2013. Detection of significantly differentially methylated regions in targeted bisulfite sequencing data. Bioinformatics 29, 1647–1653.

Huang, Q. and Dom, B. 1995. Quantitative methods of evaluating image segmentation. In Image Processing, 1995. Proceedings., International Conference on, volume 3, pages 53–56. IEEE.

Jühling, F., Kretzmer, H., Bernhart, S. H., Otto, C., Stadler, P. F., and Hoffmann, S. 2016. metilene: Fast and sensitive calling of differentially methylated regions from bisulfite sequencing data. Genome research 26, 256–262.

Kawakatsu, T., Nery, J. R., Castanon, R., and Ecker, J. R. 2017. Dynamic dna methylation reconfiguration during seed development and germination. Genome biology 18, 171.

Leenen, F. A., Muller, C. P., and Turner, J. D. 2016. Dna methylation: conducting the orchestra from exposure to phenotype? Clinical epigenetics 8, 92.

Olshen, A. B., Venkatraman, E., Lucito, R., and Wigler, M. 2004. Circular binary segmentation for the analysis of array-based dna copy number data. Biostatistics 5, 557–572.

Park, Y. and Wu, H. 2016. Differential methylation analysis for bs-seq data under general experimental design. Bioinformatics 32, 1446–1453.

Pont-Tuset, J. and Marques, F. 2016. Supervised evaluation of image segmentation and object proposal techniques. IEEE transactions on pattern analysis and machine intelligence 38, 1465–1478.

Shafi, A., Mitrea, C., Nguyen, T., and Draghici, S. 2017. A survey of the approaches for identifying differential methylation using bisulfite sequencing data. Briefings in bioinformatics.

Stouffer, S. A., Suchman, E. A., DeVinney, L. C., Star, S. A., and Williams Jr, R. M. 1949. The american soldier: Adjustment during army life. (studies in social psychology in world war ii), vol. 1.

Sun, S. and Yu, X. 2016. Hmm-fisher: identifying differential methylation using a hidden markov model and fishers exact test. Statistical applications in genetics and molecular biology 15, 55–67.

Teschendorff, A. E. and Relton, C. L. 2018. Statistical and integrative system-level analysis of dna methylation data. Nature Reviews Genetics 19, 129.

Whitlock, M. C. 2005. Combining probability from independent tests: the weighted z-method is superior to fisher’s approach. Journal of evolutionary biology 18, 1368–1373.

Wu, H., Xu, T., Feng, H., Chen, L., Li, B., Yao, B., Qin, Z., Jin, P., and Conneely, K. N. 2015. Detection of differentially methylated regions from whole-genome bisulfite sequencing data without replicates. Nucleic acids research 43, e141–e141.

Yu, X. and Sun, S. 2016a. Comparing five statistical methods of differential methylation identification using bisulfite sequencing data. Statistical applications in genetics and molecular biology 15, 173–191.

Yu, X. and Sun, S. 2016b. Hmm-dm: identifying differentially methylated regions using a hidden markov model. Statistical applications in genetics and molecular biology 15, 69–81.

